# Red squirrel territorial vocalizations deter intrusions by conspecific rivals

**DOI:** 10.1101/448894

**Authors:** Erin Siracusa, Marina Morandini, Stan Boutin, Murray M. Humphries, Ben Dantzer, Jeffrey Lane, Andrew G. McAdam

**Author notes:** Correspondence: E. Siracusa, Department of Integrative Biology, University of Guelph, 50 Stone Road East, Guelph, ON N1G 2W1, Canada *E-mail address.

## Abstract

In many species, territory defense is thought to be one of the primary functions of acoustic communication. North American red squirrels are a territorial species in which ‘rattles’ have long been thought to be the principal signal communicating territory ownership. These vocalizations have been assumed to deter intruders, thus reducing energy costs and the risk of injury associated with direct aggressive interactions. However, this hypothesis has not been directly tested. Here we used a speaker occupation experiment to test whether red squirrel rattles function to deter conspecific rivals. We studied 29 male squirrels and removed each individual from his territory twice in a paired design. During the experimental treatment we simulated the owner’s presence after its removal by broadcasting the owner’s rattle from a loudspeaker at the center of the territory once every seven minutes. During the control treatment the territory was left in silence after the temporary removal of the owner. We found that the presence of a speaker replacement reduced the probability of intrusion by 34% and increased the latency to first intrusion by 7%, providing support for the hypothesis that rattles play an active role in reducing intrusion risk. However, intrusions were not completely averted by the speaker replacement, indicating that vocalizations alone are not sufficient without other cues of the territory owner.

## Introduction

Vocal communication is thought to have several principle functions, including territorial defence (Catchpole, 1982; Catchpole & Slater, 1995; Bradbury & Vehrencamp, 2011). While the role of vocalizations in repelling conspecific rivals is typically well accepted, this function has rarely been directly demonstrated. Evidence supporting the functionality of vocalizations as a deterrent for intruders has come mostly through indirect means, via observational and correlational studies in the field, rather than direct experimental tests of functionality. For example, vocalizations commonly observed in association with intrusion events or aggressive interactions among individuals (Smith, 1978; Catchpole, 1983; Kramer & Lemon, 1983; Sharpe & Goldingay, 2009), containing characteristics such as low frequency and high intensity (believed to be associated with aggression; (Morton, 1977; Anderson & Barclay, 1995), or seasonally associated with times of important territorial defence (Catchpole, 1973; Penteriani, 2002) have typically been ascribed a territorial function. However, while suggestive, these correlative studies lack the causal evidence to support the putative defensive functionality of vocalizations.

The use of experimental playbacks is one technique that has been employed to study the role of vocalizations in territory defence. Playbacks have been used to simulate the intrusion of a rival individual by broadcasting a vocalization from a loudspeaker placed on a focal territory (Weeden & Falls, 1959). The aggressive reaction of territory owners in response to simulated intrusions has been used as evidence to support the conclusion that vocalizations play a role in territory defence against conspecific rivals. This type of aggressive response has been observed in a variety of taxa including anurans (Wells, 1977; Bastos et al., 2011; Morais et al., 2015), fishes (review: Bass & McKibben, 2003), birds (Odom & Mennill, 2010; Brumm et al., 2011; Cain & Langmore, 2015) and mammals (Barlow & Jones, 1997; Reby et al., 1999; Hayes et al., 2004; Darden & Dabelsteen, 2008). By inducing an aggressive reaction in territory owners, the use of playbacks can effectively demonstrate that vocalizations function in immediate territorial confrontations. However, by measuring the response of owners, rather than intruders, these studies fail to clarify whether vocalizations induce avoidance and function to keep conspecifics off the territory, even when confrontations are not imminent.

Muting and speaker occupation are two experimental designs that have been used in songbirds to test the hypothesis that acoustic signals function to deter territory intrusions. In muting experiments, territory owners are rendered silent via devocalizing surgical procedures (Peek, 1972; Smith, 1979). These experiments have provided empirical evidence for the territorial function of song by demonstrating that muted males experience higher intrusion rates and increased territory loss relative to controls whose ability to sing is left intact (Peek, 1972; Smith, 1979; McDonald, 1989; Westcott, 1992). In speaker occupation experiments, territory residents are removed and replaced by speakers broadcasting the owner’s song (Krebs, 1977; Krebs et al., 1978; Yasukawa, 1981; Falls, 1988; Yasukawa, 1990; Nowicki et al., 1998). In most cases, territories with a speaker replacement remain unoccupied longer and experience lower rates of intrusion than territories that are left silent, suggesting that song may be important in helping to repel intruders (Krebs, 1977; Krebs et al., 1978; Falls, 1988; Nowicki et al., 1998).

While birdsong is one of the most well-studied phenomena in animal communication, fewer studies have attempted to experimentally demonstrate the territorial function of vocalizations in other taxa. Speaker occupation experiments have been used in bicolor damselfish (*Pomacentrus partitus*; (Myrberg, 1997) and painted gobies (*Pomatoschistus pictus*; (Pereira et al., 2013), and muting experiments more recently in Lusitanian toadfish (*Halobatrachus didactylus*; (Conti et al., 2015) to show that vocalizations serve as a “keep-out” signal to other conspecifics. Due to the limitations of finding species amenable to such experimental designs our understanding of the territorial function of vocalizations in mammals has been limited to observational field studies or playback experiments that induce an aggressive response in the territory holder (Smith, 1978, Grinnell et al., 1995, Barlow and Jones, 1997, Reby et al., 1999, Grinnell and McComb, 2001, Sharpe and Goldingay, 2009). Harrington and Mech (1979) did demonstrate that simulated howling resulted in retreat or avoidance by neighbouring wolf packs, suggesting that howling serves to deter intruders and maintain territorial boundaries without direct aggression.

To experimentally test the function of vocalizations for territorial defence in a mammalian species we used a territorial tree squirrel (*Tamiasciurus hudsonicus*). North American red squirrels are small, arboreal squirrels in which both sexes defend exclusive, individual territories throughout the year. The core of each territory is a larder hoard of food resources called a “midden”(Smith, 1968). Red squirrels produce several vocalizations, of which the “rattle” is believed to be the most important for territorial defence (Smith, 1978). Rattles, unlike the vocalizations of songbirds, are not known to be associated with mating and are used by both sexes. Rattles are known to have a repeatable acoustic structure that allows for individual identification and discrimination by conspecifics (Digweed et al., 2012; Wilson et al., 2015). Smith (1978) observed that red squirrels produce rattles when another squirrel enters its territory, but also periodically when there is no apparent threat. Rattles were also observed to elicit fleeing behaviour from the intruder (Smith, 1978). This suggests that rattles may function as an advertisement of occupancy and help to maintain the spacing of individuals while minimizing direct aggressive interactions (Smith, 1978; Lair, 1990). By enabling the avoidance of aggressive interactions such as chases or fights, rattles may reduce energy costs and risk of injury (Wilson, 1975). The use of playback experiments has demonstrated that red squirrels can differentiate between neighbours and strangers as well as kin and non-kin using rattles, and that they respond more aggressively toward simulated intrusions from strangers (Price et al., 1990) and non-kin (Wilson et al., 2015). In another study, increased population density was found to increase vigilance and rattling rates in red squirrels (Shonfield et al., 2012; Dantzer et al., 2012). However, intruder pressure at high densities did not increase, which suggests that increasing rattling rates may be effective in deterring territorial intrusions (Dantzer et al., 2012). These observations support the idea that in red squirrels, rattles serve to advertise the owner’s presence to other conspecifics, maintain territory boundaries, and deter intruders. However, to date there is no direct experimental evidence to support this perceived functionality.

The aim of our study was to experimentally test whether red squirrel rattling functions to deter intruders. To assess this we employed a speaker occupation experiment and temporarily removed 29 squirrels from their territories in a paired design. During the control treatment the territory was left in silence, while during the experimental treatment we simulated the owner’s presence by broadcasting the owner’s rattle from a loudspeaker at the centre of the territory. We predicted that if rattles function to deter intruders in the absence of the territory owner then, compared to territories left in silence, territories with a speaker replacement would have: 1) a lower probability of intrusion and, 2) a longer latency to intrusion.

## Material and methods

### Population and Study Area

We studied a wild population of North American red squirrels (*Tamiasciurus hudsonicus*) in the southwest Yukon, Canada (61° N, 138° W), near Kluane National Park. The habitat of the study area is open boreal forest dominated by white spruce (*Picea glauca*; Berteaux & Boutin 2000; Humphries & Boutin, 2000; Krebs et al., 2001). This population has been monitored continuously since 1987 as part of the Kluane Red Squirrel Project (McAdam et al., 2007), on up to six study sites. We conducted our experiment on two study sites; one site was maintained as a control (40 ha) while the other study site (45 ha) was an experimental food-add site that has been supplemented with peanut butter between October and May every year since 2004 as part of a larger on-going study (Dantzer et al., 2013).

Each year in May and August we enumerated all individuals in the population and determined territory ownership using live-trapping methods and behavioural observations. We permanently tagged squirrels with uniquely numbered metal ear tags (National Band and Tag, Newport, KY, U.S.A) around 25 days old in their natal nest. Each squirrel was also given a unique combination of coloured wires that were threaded through the ear tags to allow individuals to be identified from a distance (see Berteaux & Boutin, 2000; McAdam et al., 2007 for a detailed description of study sites and project protocols).

### Rattle Recordings

Between June and August 2015 we recorded 240 rattles from 29 male squirrels (minimum 4-5 rattles each squirrel) using a Marantz^®^ Professional Solid State Recorder (model PMD660; 44.1 kHz sampling rate; 16-bit accuracy; WAVE format) with a Sennheiser^®^ shotgun microphone (model ME66 with K6 power supply; 40-20000 Hz frequency response (± 2.5 dB); super-cardioid polar pattern; Wilson et al., 2015, Shonfield et al., 2016). We collected all rattles opportunistically in the morning, between 0730 and 1100 hours. Squirrels were followed at a distance when attempting to collect a rattle and were not stimulated with a playback or otherwise provoked during rattle collection (Wilson et al., 2015). Although we cannot exclude the possibility that the observer’s presence elicited the rattles, squirrels on our study sites were well habituated to human observers. We edited the recorded rattles using Avisoft-SASLab Pro (Avisoft Bioacustics). To preserve rattle characteristics, we imported the recorded calls into Avisoft as uncompressed 16-bit .wav files. For each squirrel, we chose three rattles with minimum background noise. We adjusted rattles to ensure that all recordings had similar power (dB). We stored the three rattles, interspersed with 7-minute intervals of silence, as .wav files.

### Speaker Occupation Experiment

Between August and September 2015, we conducted temporary removals of 29 male squirrels from their territories, 15 from the control site and 14 from the experimental food addition site (*n* = 29). We trapped squirrels with Tomahawk live traps (Tomahawk Live Trap Co., Tomahawk, Wisconsin), and removed each squirrel from his territory twice, once as a treatment and once as a control (order of treatment and control was randomly assigned). Treatments were conducted no less than 3 days apart (range: 4-41 days). Once trapped, we placed the squirrel in a modified box (41 cm x 17.5 cm x 19 cm) to help keep the squirrel quiet and calm for the duration of the removal (Donald & Boutin, 2011). A small amount of peanut butter and some spruce cones were also provided in the modified box. We then moved the squirrel 20-30 m away from his midden, and placed the individual in a shady location. Care was taken not to place the removed individual on another squirrel’s territory.

Each trial commenced immediately following the removal of the territory owner. When removed as a control, we left the territory owner’s midden silent. When removed as a treatment, we placed a Saul Mineroff SME-AFS field speaker with a playback range of 0.1-22.5 kHz face-up on the ground at the centre of the squirrel’s midden. We then broadcast the owner’s rattle from this speaker at a level between 65-75 dB (Shonfield et al., 2016) at 7-minute intervals (Dantzer et al., 2012) for the duration of the removal. We checked the power of each recording in the field, prior to the start of the speaker replacement, using a digital sound level meter measured 2 m from the upwards-facing speaker.

During each removal an observer monitored the midden from a distance of no less than 5 m away from the midden center. If an intrusion was witnessed the observer recorded the time of intrusion and the identity of the intruding squirrel. An intrusion was defined as occurring when a squirrel moved onto the pile of cone scales, which is considered to define the edges of a squirrel’s midden. Each removal lasted 120 minutes or until an intruder arrived, whichever came first, after which point we released the captive squirrel at the site of capture. This research was approved by the University of Guelph Animal Care Committee.

### Statistical analysis

To assess whether the presence of a speaker replacement reduced the probability of a squirrel intruding and the latency to intrusion we used a Cox proportional hazard mixed effects model (PHMM) in the R package coxme (version 2.2-5: Therneau, 2015). We used a survival analysis approach because removals in which no intruder was observed in the 120 min time-period had censored values for time to intrusion. The response variable for Cox proportional hazard models is a single measure called the hazard of intrusion. This response variable takes into account both the probability of intrusion (1 = yes there was an intrusion, 0 = no intrusion observed) as well as the time to intrusion (measured in minutes). For the purposes of interpretability, hereafter we will refer to the outcome of this model as the ‘risk of intrusion’. In all cases, a high risk of intrusion is equivalent to a high probability of intrusion and a fast time to intrusion.

Our model included speaker treatment (speaker or no speaker) and study site as categorical predictors. We also included squirrel identity as a random intercept term to account for our matched pairs design. Our PHMM was found to sufficiently meet the proportional hazards assumption. We examined dfbeta residuals to ensure that there were no influential observations. All analysis were performed using R software 3.3.2 (R Core Team, 2016). For the following results, we considered differences statistically significant at α=0.05 and report all means ± SE, unless otherwise stated.

## Results

Our Cox proportional hazards model revealed that the risk of intrusion was significantly lower in the presence of the speaker replacement, meaning that playbacks of the owner’s rattle reduced the overall probability of intrusion and also delayed the time to intrusion by a neighbouring squirrel (β = -0.92 ± 0.33, *z* = -2.77, *P* = 0.01). There were no differences in the risk of intrusion between the two study sites (β = -0.24 ± 0.32, *z* = -0.75, *P* = 0.45). For each treatment we conducted 29 temporary removals of territory owners. Twenty-five intrusions occurred when no speaker replacement was present (86% of removals) and only 15 intrusions were observed when a speaker was used to broadcast the rattle of the territory owner (52% of removals). The proportion of intrusions with the speaker replacement was significantly lower than when the territory was left silent (McNemar’s Chi-squared test; χ^2^ = 5.79, df = 1, *P* = 0.02; Figure 1). However, the average time to first intrusion on empty territories was 52.20 minutes (SD: ± 36.84 min; range: 0-120 min) and on speaker occupied territories was 60.13 minutes (SD: ± 38.62 min; range: 15-120 min), which were not significantly different (Welch Two Sample t-test; *t* = -0.64, df = 28.50, *P* = 0.74; Figure 2). Thus, over a 120-minute period the presence of a rattle vocalization reduced the probability of intrusion by 34% but only reduced the time to intrusion by 7%, suggesting that the reduced risk of intrusion was primarily due to a reduced probability of intrusion.

**Figure 1.**
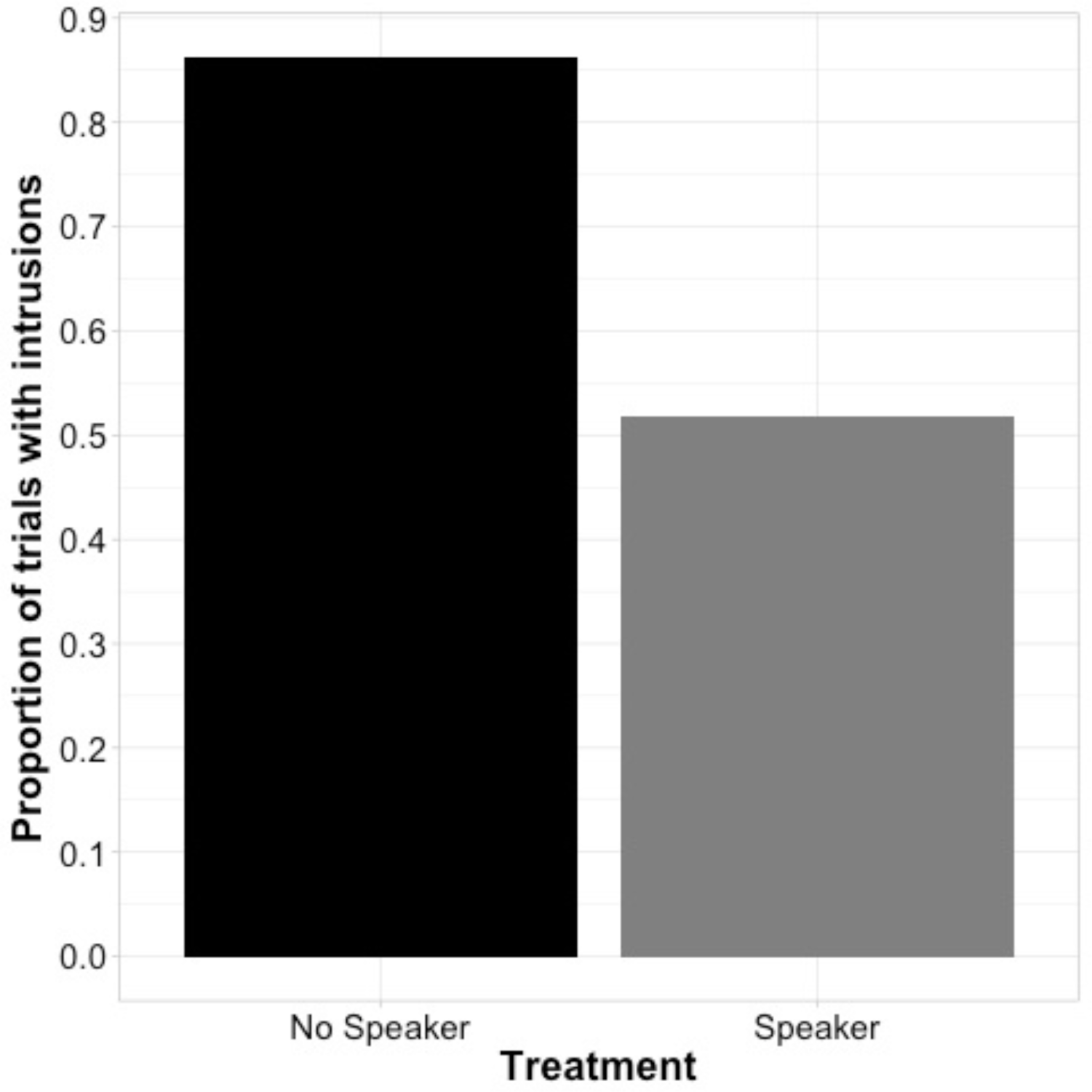
The effect of a speaker replacement following the temporary removal of a territory owner on the proportion of trials with intrusions (McNemar’s Chi-squared test; χ^2^ = 5.79, df = 1, *P* = 0.02; *n* = 29).

**Figure 2.**
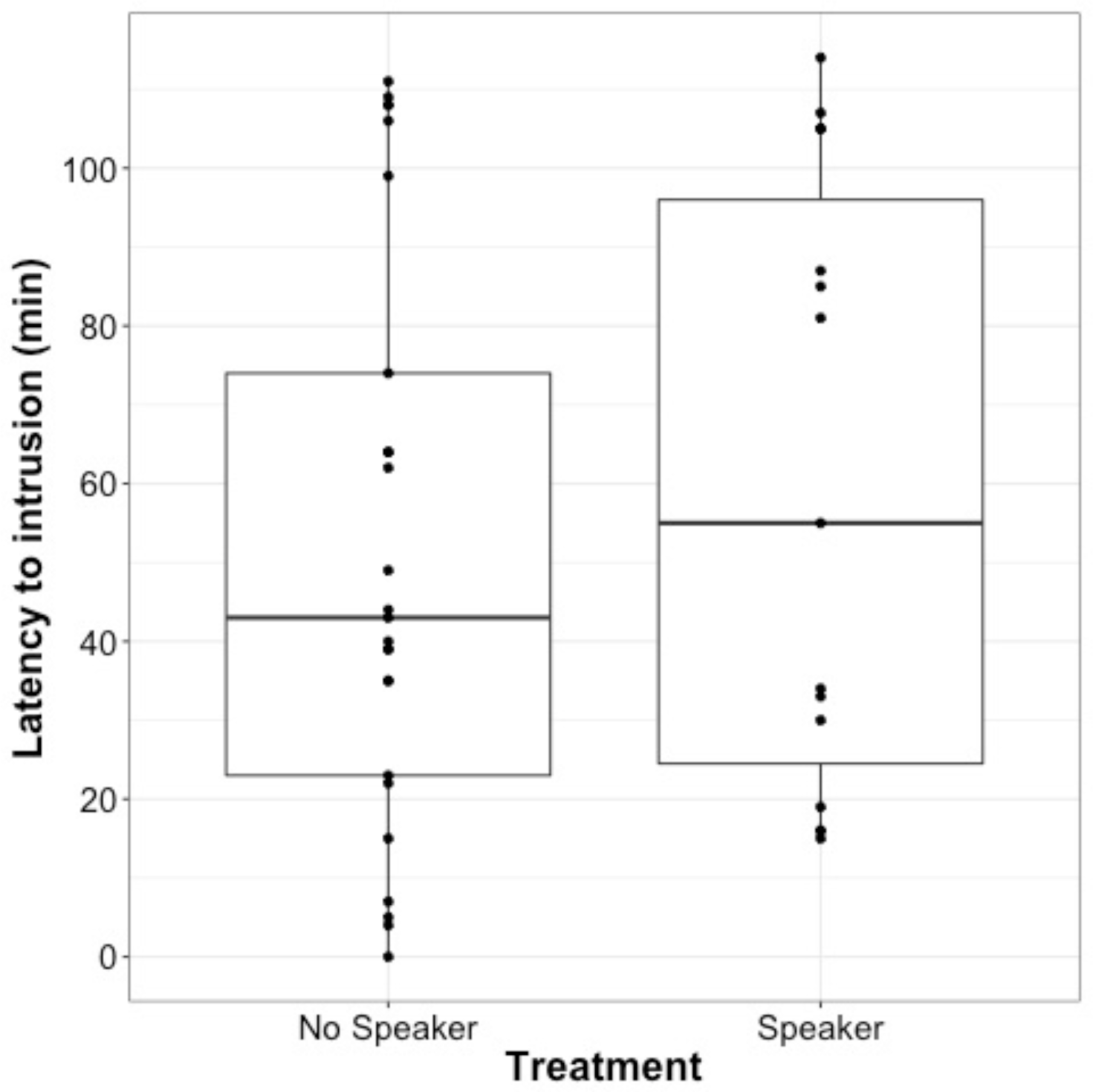
The effect of a speaker replacement following the temporary removal of a territory owner on the latency to intrusion (Welch Two Sample t-test; *t*= -0.64, df = 28.50, *P* = 0.74; *n* = 29).

## Discussion

In this study we investigated the territorial function of vocalizations in the North American red squirrel. Specifically, we tested the role of rattles in discouraging conspecific intruders. The results from this study confirmed our hypothesis that red squirrel rattles reduce intrusion risk by demonstrating that territories occupied by a speaker replacement had a significantly lower probability of intrusion and a delayed time to first intrusion compared to territories that were left empty.

Our results, obtained using a speaker occupation design, are the first to experimentally demonstrate the territorial function of vocalizations in a mammal. Moreover, our findings support the conclusions of previous speaker occupation studies in both songbirds and territorial fishes that have demonstrated the importance of acoustic signalling for territory defence and retention (Krebs, 1977; Krebs et al., 1978; Falls, 1988; Myrberg, 1997; Nowicki et al., 1998; Pereira et al., 2013). However, while in red squirrels the number of intrusions was clearly reduced by the rattle playback, the presence of the speaker replacement alone was not 100% effective in deterring intrusions. Of the 29 territories which received a speaker playback, 15 experienced a territorial intrusion, which is just over half of the territories (52%), suggesting that visual cues, or other stimuli from the territory owner, are also an important component of territory defence. This is consistent with previous speaker occupation studies which demonstrate lower intrusion rates or delayed intrusions in the presence of playbacks, but do not preclude intrusions altogether (Falls, 1988; Myrberg, 1997; Nowicki et al., 1998). Falls (1988) suggested that the effectiveness of the speaker occupation design may depend in part on the conspicuousness of the territorial species that is being studied. Speaker replacement should be most effective in species that are visually inconspicuous because it allows the façade of occupancy to be maintained for a longer period of time. Red squirrels are usually fairly visible to their neighbours as they are diurnal and spend a significant portion of their time foraging and feeding on or around their territory. Red squirrels will also actively chase intruders off their midden if necessary, although such direct acts of aggression are usually rare (Gorrell et al., 2010; Dantzer et al., 2012).

Given the potential importance of the physical presence of the territory owner, it is not surprising that the speaker replacement experiments were not wholly effective in deterring red squirrel intruders. In fact, in some studies, speaker replacements initially induced avoidance of ‘occupied territories’, but had unexpected effects on intrusion rates. For example, in red-winged blackbirds there were reduced rates of fly-through, but not neighbour trespass, on speaker replacement territories, suggesting that visual displays may be crucial for territory retention (Yasukawa, 1981). In painted gobies the presence of a playback ultimately led to higher rates of territory intrusion than in territories that were left silent. Painted gobies approached ‘occupied’ nests more frequently than silent nests but then demonstrated avoidance after approaching the agonistic calls. However, when the gobies were unable to associate the sound with the physical presence of a territory holder they proceeded to intrude on the ‘occupied’ territory (Pereira et al., 2013). Anecdotally, we observed similar patterns of behaviour in red squirrels. Potential intruders that appeared at the edge of the territory initially appeared to be deterred by the broadcast rattle, however, after remaining on the territory for several minutes without sighting the territory owner, these loitering individuals entered the midden and, when not chased off by a resident squirrel, proceeded to pilfer food resources. Without the visual presence of the territory owner, potential intruders appear to become habituated to a repetitive acoustic stimulus. While we have confirmed the significance of rattles in red squirrel territory defence, altogether these results suggest that, similar to other acoustic signals, rattles only serve as a temporary deterrent to territorial intrusions until the absence of the territory owner is confirmed. This underscores the importance of the combined effects of visual and acoustic cues for deterring conspecific rivals.

A more precise understanding of how rattles function in territory defence is still lacking. Red squirrel rattles are known to have unique signatures that allow for individual identification by other conspecifics (Wilson et al., 2015), but it is unclear if rattles will indiscriminately discourage intruders regardless of whether that rattle is from a resident or non-resident conspecific. Yasukawa (1981) suggested that one reason the speaker replacement did not deter neighbour trespass in red-winged blackbirds is because song types were used which were unfamiliar to neighbouring conspecifics. Yasukawa speculated that neighbouring males interpreted this as an attempted territory establishment by a new individual and therefore responded by either attempting to re- negotiate boundaries or claim the abandoned territory before the new male had time to fully settle (Yasukawa, 1981). Here we used only the calls of the territory owners in our speaker replacements. Future studies should employ the use of speaker occupation experiments using calls from different signallers to better understand if red squirrels are able gather and respond to more complex information about their social environment.

## Acknowledgements

We would like to thank the Champagne and Aishihik First Nations on whose land the study sites for this project were located. In particular, we would like to thank Agnes MacDonald and her family for long-term access to her trapline. A special thank you to J. Robertson and S. Sonnega for assistance with field data collection. Research support was provided by Grants-in-Aid of research from the American Society of Mammalogists and the Arctic Institute of North America (E. Siracusa), as well as funding from the Natural Sciences and Engineering Research Council of Canada (A.G. McAdam).

